# Monocyte and Lymphoid Lineages Share a Common Precursor Distinct from Neutrophils in Hematopoiesis

**DOI:** 10.64898/2025.12.25.696474

**Authors:** Yongjian Yue, Sujin Chen

## Abstract

During lineage commitment in hematopoiesis, hematopoietic stem cells (HSCs) differentiate into distinct lymphoid and myeloid lineages. Within the myeloid lineages, granulocytes and monocytes have been considered to originate from granulocyte-macrophage progenitors (GMPs) or common myeloid progenitor (CMPs). Single-cell sequencing has refined this hematopoietic hierarchy, revealing that neutrophils and basophils diverge early during commitment. Although current hierarchy models propose that neutrophils and monocytes arise from common neutrophil-macrophage progenitor (NMP) or GMP, their early differentiation trajectories remain inconsistent, leaving the origin relationship between neutrophils and monocytes unclear. This study aimed to clarify the developmental origin of neutrophils and monocytes during hematopoiesis. Leveraging a previously established single-cell atlas of hematopoietic progenitor cells (HPCs), we refined a trilineage hierarchy model, revealing the direct commitment of neutrophils from early multipotent progenitors (MPCs) and their divergence from lymphoid lineages. HOPX was identified as a key regulator during lymphoid lineage commitment. Using HOPX-based lineage tracing, we reconstructed the differentiation trajectory of neutrophils and monocytes. Lineage-tracing results revealed that HOPX-derived tdTomato positive cells contribute extensively to lymphoid lineages and monocytes, but not to neutrophils or erythroid cells. Ex vivo primary macrophage cultures derived from bone marrow cells yielded consistent findings. Our results indicate that monocytes are committed from same precursor with lymphoid lineages, and diverge from neutrophils. Moreover, we propose that classical common lymphoid progenitors (CLPs), redefined as lymphoid-monocyte progenitors (LMPs), rather than GMPs, constitute the likely ancestral source of monocyte lineage. These findings challenge the dogma of hematopoietic hierarchy and open new avenues for understanding lineage specification. Future studies involving conditional knockout of HOPX will provide further mechanistic validation of these conclusions.

## Introduction

Immunology cell lineages originate from HSCs during hematopoiesis, a process described by the classical lymphoid-myeloid dichotomy (1). HSCs, a rare population in the peripheral blood, possess the capacity to generate all lineages with tree-like patterns in classical hematopoietic hierarchical models (2). Granulocyte (including neutrophil, basophils, and eosinophils) and monocytes (macrophage and dendritic cells) have long been considered to originate from common precursors of common myeloid progenitors (CMPs) or granulocyte-monocyte progenitors (GMPs).

With the advantages of single cell sequencing, discrete hematopoietic hierarchy was constructed that showed diverse differentiation trajectories during lineage commitment (3, 4). Distinct multipotent hematopoietic progenitor cells (HPCs) were proposed, such as transitional progenitors of LT-HSC, ST-HSC, LMPP, MPP1, and MPP2 (5). Inversely, a continuum model of hematopoiesis has been proposed (6, 7), suggesting that there are no strict boundaries between the different hierarchical levels of HPCs. Transitional progenitor states are not homogeneous populations, and unilineage-restricted cells can emerge directly from a continuum of low-primed undifferentiated HSCs (8). However, the inconsistent definitions of HPCs, lineage heterogeneity, and dynamic differentiation impede the characterization of lineages. The unclear delineation of transitional states during HPC differentiation caused controversial definitions of CMPs, GMPs, and lymphoid-primed multipotent progenitors (LMPPs) (9). At least three distinct differentiation trajectories have benn proposed for granulocyte and monocyte commitment (3). One emerging model suggests that granulocyte and monocyte lineages originate from NMPs, which derived from the same LMPP precursors as CLPs (9). Further studies indicated that basophils and eosinophils were derived from same precursor with megakaryocytic–erythroid lineages (9, 10), implying that neutrophils may follow an independent path from other granulocytes during early lineage commitment. Although different hierarchical models assign neutrophil and monocyte origins to distinct early precursors (e.g., CMP), there is consensus that they arise from common progenitors, such as GMPs or NMPs. However, additional complexity is introduced by studies showing that dendritic cells can develop not only from monocytes but also from lymphoid progenitors, though findings remain conflicting (11-13). Consequently, the precise origin, differentiation trajectories, and fate decisions governing neutrophils and monocytes in hematopoiesis remain ambiguous..

Our previous study established a revised trilineage priming model of hematopoietic hierarchy that demonstrated neutrophil progenitors (NMPs) were primed simultaneously with GAPs and CLPs during the initial bifurcation of hematopoiesis (4). This early tri-lineage priming is orchestrated by the key regulators *GATA2, HOPX*, and *CSF3R*. However, the cellular origin of monocytes (which give rise to macrophages and dendritic cells) remained unresolved. In this study, we employed a conditional HOPX lineage-tracing mouse model to elucidate the developmental relationship among neutrophils and monocytes. Our results may reveal the characterization of neutrophil and monocyte lineages commitment, facilitating advancements in immunity modulation and therapeutic strategies.

## Methods

### Single cell RNA sequencing atlas of HPCs

Single cell RNA sequencing atlas of HPCs was constructed in previous study(4). Expression profiles were analyzed by Seurat. Pseudotime trajectory analysis was performed with Monocle 3.

### Lineage tracing in a Hopx-based genetic model

To specifically label and trace Hopx-expressing progenitor and differentiated cells, we generated lineage-tracing mice by crossing Hopx-CreERT2 animals (GemPharmatech Co., Ltd) with Rosa26-LSL-tdTomato reporter mice (Cyagen Biosciences Inc). The resulting double-transgenic Hopx-CreERT2; Rosa26-LSL-tdTomato mice were housed in the Animal Center of Shenzhen People’s Hospital, and all procedures were approved by the Institutional Animal Care and Use Committee of the hospital. For Cre-mediated recombination, tamoxifen (Sigma, T5648) was dissolved in corn oil (Sigma, C8267) at 20 mg/mL, vortexed gently, and incubated at 37 °C for 1 h. Eight- to ten-week-old mice received intraperitoneal injections of tamoxifen (100 mg/kg body weight) daily for five consecutive days. Control animals were injected with corn oil only.

### Flow cytomerty

Following tamoxifen-induced Cre recombinase activition over five consecutive days, the mice were euthanized. Bone marrow and peripheral blood cells were harvested. Whole peripheral blood was lysed by ACK buffer for 3 minutes. The resulting cell suspensions were stained with antibodies against CD3-FITC, NK1.1-APC, CD19-PC5.5, CD11b-PC7, GR1-APC, and TER119-FITC (Biolegend). Flow cytometry analysis was conducted to identify tdTomato labeled cell populations.

### In vitro culture of tamoxifen-induced bone marrow cells

Bone marrow cells were havested after five consecutive days of tamoxifen induction. Erythrocytes were removed by lysis in ACK buffer for 3 minutes. Primary cells were cultured in RPMI-1640 medium supplemented with 10% FBS. For monocyte cell differentiation, cytokine factors of M-CSF (20ng/ml) and G-CSF (20ng/ml) were added to the culture medium. Surface expression of CD11b-PC7 and F4/80-FITC (Biolegend) was assessed by flow cytometry to determine cell identity and the proportion of tdTomato-positive cells.

## Results

### The ambiguous differentiation trajectories of monocyte lineage

To clarify the differentiation trajectories of monocyte lineages, it is essential to define and characterize the transitional states along monocyte progenitor differentiation. In our previous study, most HPCs were redefined by a scRNA transcriptional atlas (Figure 1A)(4). That work identified three branching lineages--- megakaryocytic–erythroid progenitors (GAP), CLPs, and NMPs---during the first stage of hematopoietic differentiation (Fig 1A)(4). However, we didn’t detect clear evidence of a monocyte differentiation trajectory, even though the CLPs and NMPs were distinctly identifined. These data confirmed that monocyte commitment was distinction from the megakaryocytic–erythroid linage, but two possibilities remain: monocyte may originate from CLPs or from NMPs (Fig 1A). We further performed pseudotime trajectory analysis by monocle3, which also revealed no apparent trajectory toward monocyte commitment (Fig 1B).

**Figure 1.**
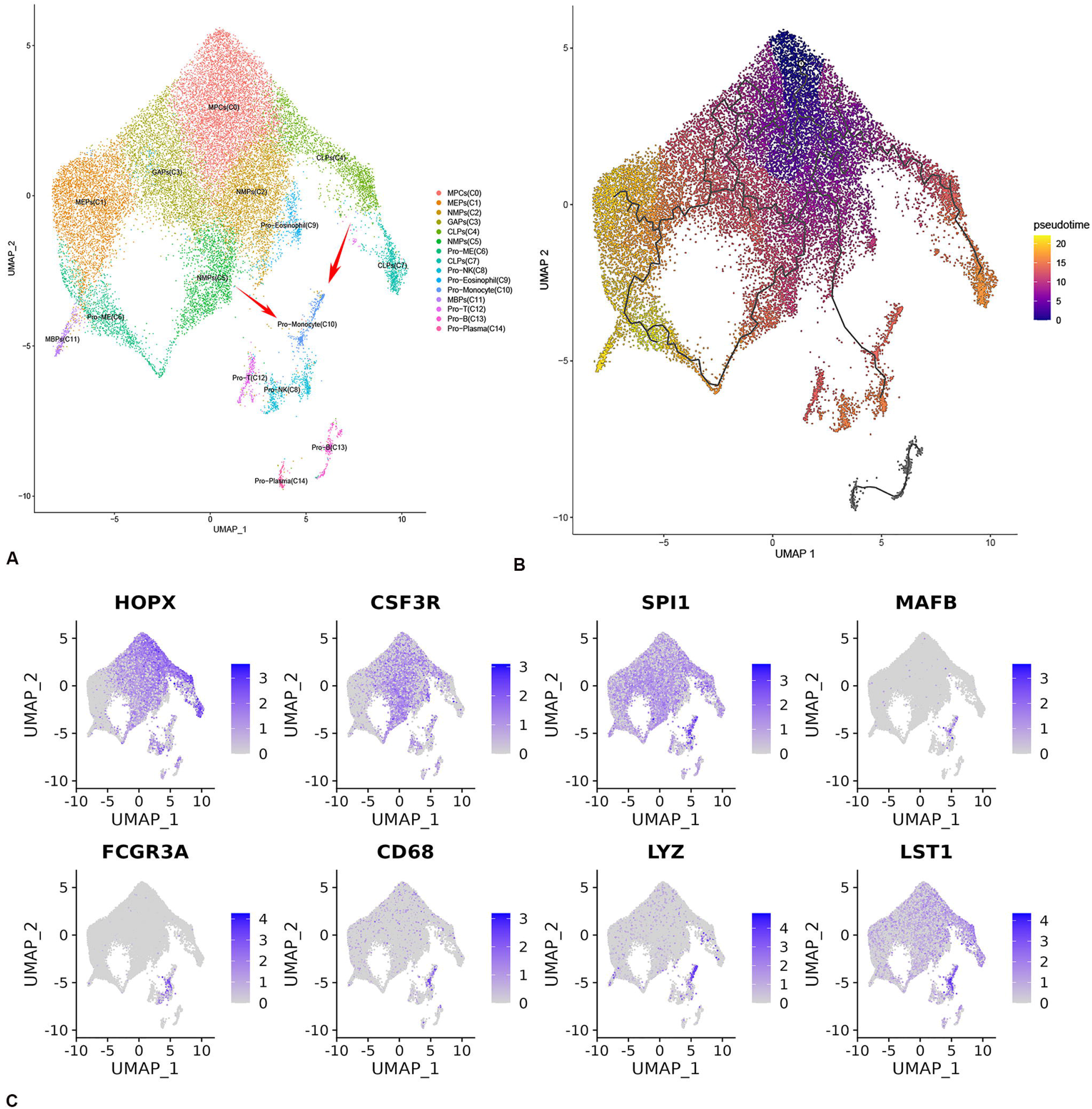
The predicted differentiation trajectories of monocyte lineages. A, T-SNE plot of redefined HPCs, depicting two potential differentiation trajectories of monocyte lineages. B, Pseudotime trajectory analysis plot by monocle 3 displaying ambiguous differentiation trajectories of monocyte subsets, including an entirely independently trajectory. C, Feature plots of CLP, NMP, and monocyte progenitor marker genes illustrating similar expression profiles of *LYZ* and *LST1* in CLP and monocyte lineages. HPC, hematopoietic progenitor cells; CLP, peripheral blood mononuclear cell; NMP, neutrophil–monocyte progenitors.

### HOPX acts as a key fate-determining factor for CLP commitment

To elucidate the developmental relationship between monocyte progenitors and other lineages, we analyzed genes co-expressed in monocyte progenitors and either CLPs or NMPs (Fig 1C). Lysozyme (LYZ) and leukocyte-specific transcript 1 (LST1) showed co-expressed in CLPs and monocyte progenitors. This similar expression profile indicated that monocytes may also originate from CLPs (Fig 1C). We also observed that the known myeloid transcription factor SPI1 was broadly expressed across most myeloid progenitors and therefore could not be serve as a specific marker for lineages discrimination.

HOPX has been widely reported as a marker gene of HSCs. In our previous model, early hematopoietic progenitor commitment bifurcates into three lineages regulated by GATA2, HOPX, and CSF3R(4). Expression analysis revealed that HOPX and CSF3R were not expressed in the GAP lineage. CSF3R was highly expressed in early priming stages and in NMPs (Fig 1C). CSF3R likely plays a role in multiple lineage-commitment processes. In contrast, HOPX was highly expressed specifically in CLPs, indicating that it serves as a distinguishing fate-determining factor for commitment toward the CLP and NMP lineages.

### Linage tracing of HOPX reveals that monocytes originate from a common precursor shared with CLPs

To validate the origin of monocyte lineage, we performed lineage tracing of HOPX using conditional Hopx-CreERT2; Rosa26-LSL-tdTomato mice. Lineage-tracing analysis showed no detectable tdTomato signal in neutrophils (Figure 2a) or erythroid cells (Figure 2b), which aligns with the established view that CLP-derived lineages are independent of neutrophil and erythroid commitment during hematopoiesis. Flow cytometry results showed a high percentage of tdTomato-positive cells among CLP-committed lineages, including T cells, B cells, and NK cells (Figure 3a, b). Furthermore, *in vitro* culture of tamoxifen-treated bone marrow cells demonstrated that a substantial proportion of induced macrophages were tdTomato-positive (Figure 3c). Fluorescent microscope confirmed tdTomato expression in macrophages (Figure 3d). Together, these results revealed that monocyte share a common precursor with CLP lineages, which we redefined as lymphoid-monocyte progenitors (LMPs).

**Figure 2.**
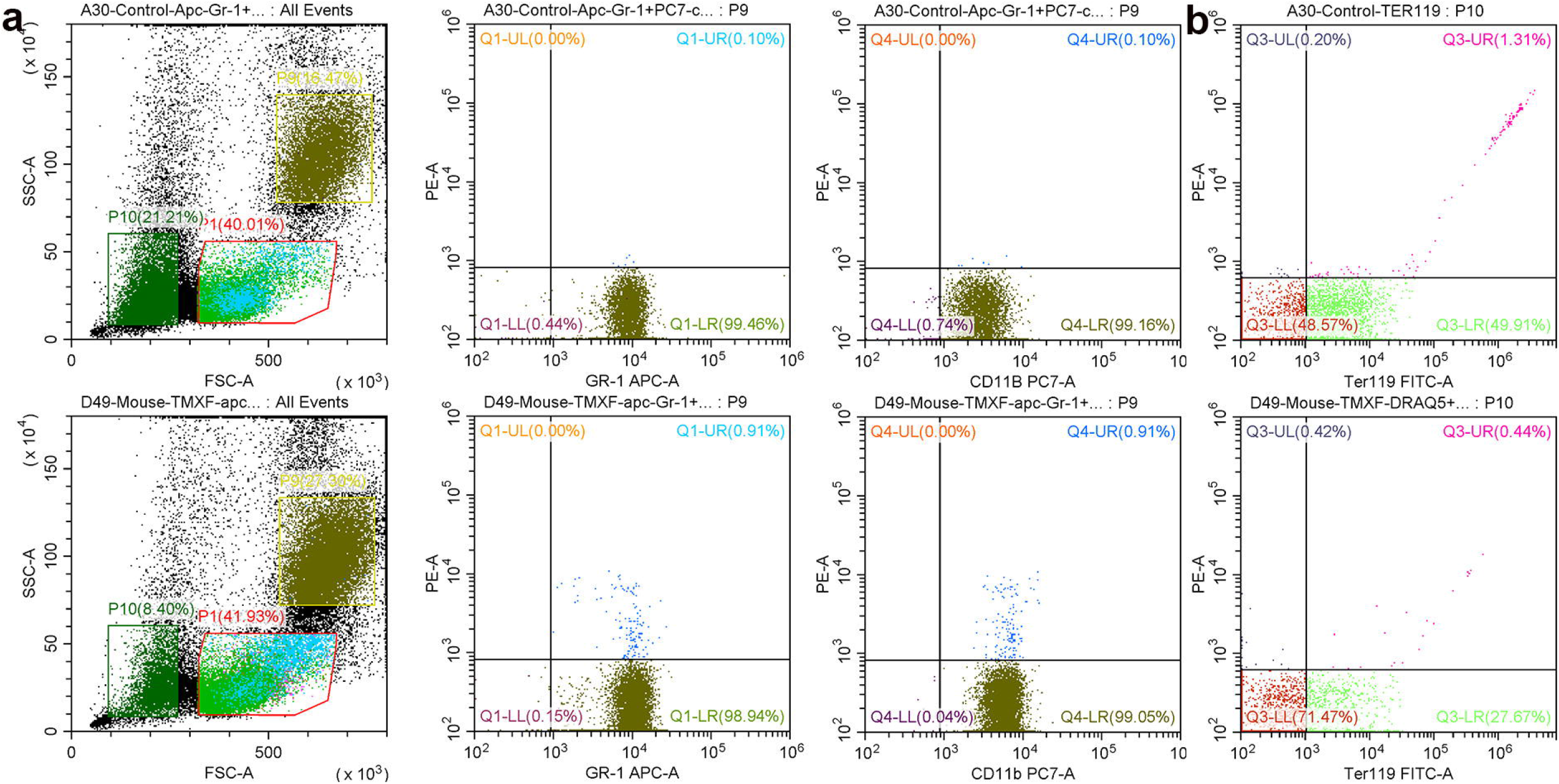
Flow cytomerty analysis of tdTomato labeling in neutrophils and erythroid cells. a, Flow cytomerty plot illustrating that neutrophils cells (Gr-1 +CD11b+) are not labeled with tdTomato fluorescence. b,Ter119-FITC positive erythroid cells are showing negative expression of tdTomato (PE).

**Figure 3.**
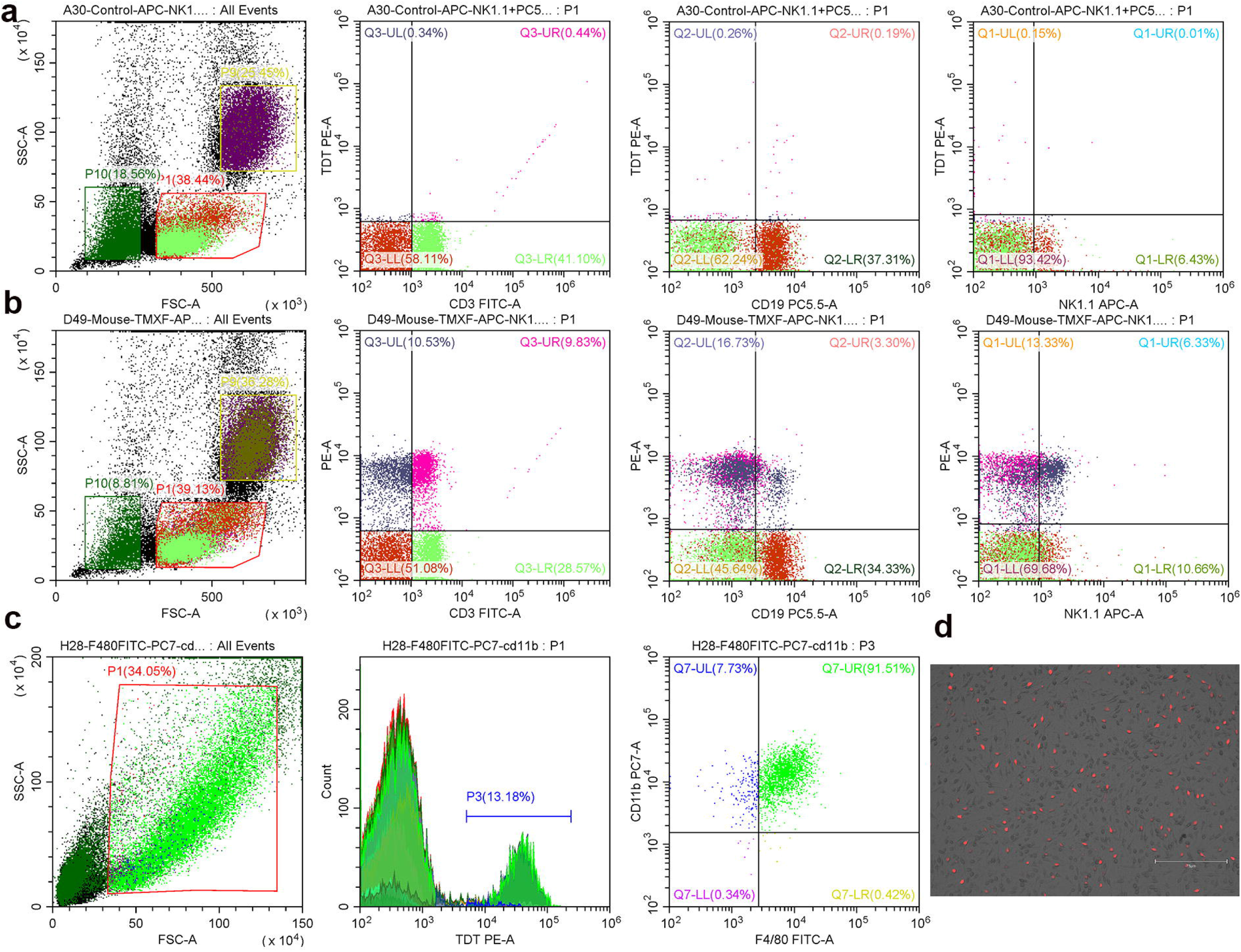
HOPX lineage tracing reveals tdTomato labeling in CLP-derived lineages and macrophages. a, In control mice, T, B, and NK cells show no tdTomato (PE) signal. b, Flow cytometry analysis demonstrating tdTomato labeling in all mature CLP-derived cell types from tamoxifen-induced mice. c, *In vitro* macrohpages culture from tamoxifen-induced bone marrow cells illustrating tdTomato labeling cells are macrohpages (CD11b+F4/80+). d, Fluorescent microscope plot showing tdTomato expression in the cultured macrophages.

### Reforming the hierarchy of monocyte lineage commitment

The conflicting definitions of CMP and GMP have traditionally relied on surface markers such as CD45RA, CD135, and CD123. Recent studies have sought to redefine transitional states, including lymphoid-primed multipotent progenitors (LMPPs), which is proposed to differentiate into lymphoid lineages, neutrophils, and macrophages, yet its definition remains inconsistent. To clarify these conflicts, we analyzed the expression profile of CD135 or CD123, and found that both markers are broadly expressed across the redefined CLP branch (Figure 4a). This indicated classical surface-markers based definitions of CMP and GMP likely capture a heterogeneous mixture of multipotent progenitors, potentially explaining how CD123-positive populations can be induced toward monocyte lineages. Leveraging the reformed hematopoietic hierarchy, we have updated the model of monocyte lineage commitment (Figure 4b). Our model shows that after the initial trilineage priming stage of hematopoiesis, NMPs commit exclusively to the neutrophilc lineage, which develops independently of other lineages. Under the regulation of HOPX, MPCs are priming into LMPs, which possess the dual potential to differentiate into both monocytic and lymphoid lineages.

**Figure 4.**
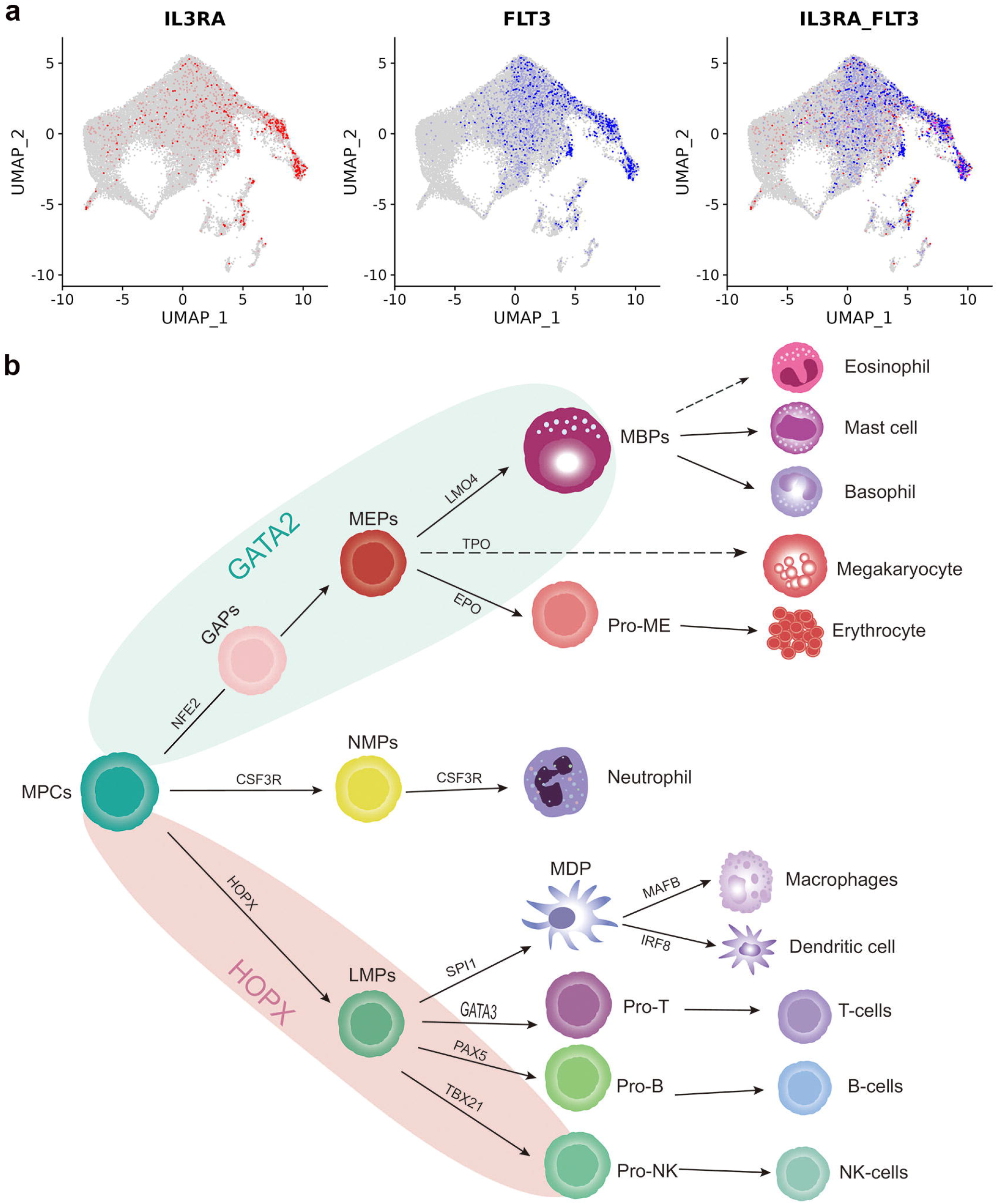
A revised hierarchy of monocyte lineage commitment. a, Feature plots are illustrating the widespread expression of CD135 and CD123 within the redefined CLP branch. B, Revised hierarchy of monocyte lineages based on HOPX lineage tracing, showing that MDPs share a common LMP with lymphoid lineages. Shaded areas are indicating GATA2 and HOPX expression domains. CLP, common lymphoid progenitors; MDP, monocyte–dendritic cell progenitors; LMP, lymphoid and monocyte progenitors.

## Discussion

Prior studies yielded conflicting hierarchy models of neutrophils and monocytes, and the differentiation trajectories of monocyte lineages remained ambiguous. Our study demonstrated that MDPs originated from progenitors distinct from those of neutrophils, but shared with lymphoid lineages. We proposed that classically defined CLPs, redefined here as LMPs, likely represented the common ancestor for both monocyte and lymphoid lineages. Consequently, our findings challenged the prevailing hematopoietic dogma that monocytes and neutrophils originated from the same precursor progenitors.

Our previous study showed that neutrophils are directly primed from initial hematopoietic progenitors and develop independently with megakaryocytic–erythroid and lymphoid lineages (4). However, the origin of monocyte-dendritic cell progenitors remained unconfirmed. MDPs are defined as a common precursor of monocytes, macrophages and DCs, which derived from CMP or GMP (14). Previous study indicated that Ly6C+ progenitors generated major monocyte subsets and macrophages, but not DCs or neutrophils (14). Moreover, GMPs and MDPs were reported to independently produce functionally distinct monocytes (15), suggesting that these progenitors possess different potentials in monocytes subtypes generation. However, these conclusions were not fully conclusive. One limitation was that the definitions and distinction between GMPs and MDPs relied heavily on a limited set of specific surface markers, such as LY6C, FLT3, and CD115. That would cause the selection of heterogeneous progenitor populations in functional assays. Collectively, these studies highlighted that GMPs and MDPs may be independently mobilized to produce specific myeloid cell types, and implied that neutrophils likely originated from distinct precursor subtypes. Recent studies further clarified that the differentiation trajectories of neutrophils differe with those of basophils and eosinophils, thereby simplifying efforts to trace the origin of neutrophils (9). Despite these advances, the precise origin and differentiation trajectories of neutrophils and monocytes within the hematopoietic hierarchy remained unclear.

Single-cell sequencing has enabled the identification and redefinition of transitional progenitor states, such as LMPP, Pre-GM, and Pre-NM (6, 16). These redefined populations formed the basis of reconstructed hierarchy models, which proposed three main differentiation trajectories for neutrophils and monocytes: a classical model deriving from CMP (shared with MEPs); and a mixed model deriving from both CMP and LMPP; and a third model deriving from the LMPP (shared with CLP) (3). Notably, all three hierarchy models suggested that neutrophils and monocytes originated from a common precursor progenitors (NMP or GMP). Inversely, the continuum hematopoietic model proposed that there were no clear boundaries between the different hierarchical levels of progenitor cells(17), implying that lineage commitment occurred from a broad progenitor pool rather than from a single, discrete precursor (6). Our previous study effectively delineated transitional progenitors and clearly distinguished the differentiation states along lineages commitment, thereby reconciling previously controversial definitions of CMP, GMP, and LMPP (4). T This refined framework for HPCs and lineage commitment reduced the confusion introduced by earlier, disparate models. Furthermore, the regulatory factor HOPX was identified as playing an essential role in monocytes and CLP commitment. Lineage tracing using HOPX revealed that neutrophils and monocytes followed distinct differentiation trajectories, with neutrophils deriving from an independent branch. This finding challenged the long-standing dogma that neutrophils and monocytes originate from common precursor progenitors.

Our lineage-tracing results showed that monocytes and lymphoid cells originated from a common redefined progenitor population, which we termed lymphoid-monocyte progenitors (LMPs). HOPX was identified as a marker of HSC(18). Our previous study showed that during the initial commitment phase, the bifurcation of the LMP lineage is mainly orchestrated by HOPX (4). Accordingly, HOPX-based lineage tracing effectively reconstructed the developmental relationship between lymphoid and monocyte differentiation. We found that HOPX was expressed not only in LMPs but also in MPCs. However, only specific lineages were ultimately labeled by HOPX-derived fluorescence, suggesting that HOPX-positive progenitor cells had already undergone fate restriction even at the MPC stage. Most conventional dendritic cell (cDC) are derived from the myeloid lineage (19). Although CLPs have been reported to possess potential for cDC differentiation, prior lineage tracing studies via CD19 or IL7R did not demonstrate CLP contribution to monocytes (11). Unlike HOPX, CD19 and IL7R are not markers of early MPCs but are expressed at low level in relatively later and restricted CLPs subsets. Thus, the common LMP precursor likely represents a multipotential subset, whereas IL7R subset appears to be fate-restricted toward the lymphoid lineage. These observations further support our conclusion that progenitor restricted occurs much earlier than previously recognized.

In conclusion, our study reconciled the previously controversial differentiation trajectories of monocyte and neutrophils. We demonstrated that monocyte lineages are committed from a common precursor (LMP) with lymphoid lineages but diverge from neutrophils. Our study provided new perspectives for the precise control of progenitor cell differentiation and advances in immunology modulation.

## Acknowledgments

This work was supported by the Shenzhen Science and Technology Program (JCYJ20240813103819026). Platform supported by a grant from the Shenzhen Clinical Research Center for Respiratory Disease (LCYSSQ20220823091203007) and Shenzhen Key Laboratory of Respiratory Diseases (SYSPG20241211173920041).

## Authors’ contributions

Y.Y. conceived the original idea and designed the study. Y.Y. conducted the bioinformatics analyses of the scRNA data. Y.Y. wrote and revised the manuscript. S.C. conducted the sample collection and animal experiment. S.C. and Y.Y. conducted the primary culture and flow cytometry detection. All authors approved the final draft of the manuscript.

## References

1. Akashi K, Traver D, Miyamoto T, Weissman IL. A clonogenic common myeloid progenitor that gives rise to all myeloid lineages. Nature. 2000;404(6774):193–7.

2. Cheng H, Zheng Z, Cheng T. New paradigms on hematopoietic stem cell differentiation. Protein & cell. 2020;11(1):34–44.

3. Jacobsen SEW, Nerlov C. Haematopoiesis in the era of advanced single-cell technologies. Nat Cell Biol. 2019;21(1):2–8.

4. Chen L, Sun Q, Li G, Huang Q, Chen S, Fu Y, et al. Redefining hematopoietic progenitor cells and reforming the hierarchy of hematopoiesis. bioRxiv. 2024:2023.01.27.524347.

5. Wilson A, Laurenti E, Oser G, van der Wath RC, Blanco-Bose W, Jaworski M, et al. Hematopoietic Stem Cells Reversibly Switch from Dormancy to Self-Renewal during Homeostasis and Repair. Cell. 2008;135(6):1118–29.

6. Karamitros D, Stoilova B, Aboukhalil Z, Hamey F, Reinisch A, Samitsch M, et al. Single-cell analysis reveals the continuum of human lympho-myeloid progenitor cells. Nature immunology. 2017;19(1):85–97.

7. Velten L, Haas SF, Raffel S, Blaszkiewicz S, Islam S, Hennig BP, et al. Human haematopoietic stem cell lineage commitment is a continuous process. Nat Cell Biol. 2017;19(4):271–81.

8. Laurenti E, Gottgens B. From haematopoietic stem cells to complex differentiation landscapes. Nature. 2018;553(7689):418–26.

9. Drissen R, Thongjuea S, Theilgaard-Mönch K, Nerlov C. Identification of two distinct pathways of human myelopoiesis. Science immunology. 2019;4(35).

10. Drissen R, Buza-Vidas N, Woll P, Thongjuea S, Gambardella A, Giustacchini A, et al. Distinct myeloid progenitor-differentiation pathways identified through single-cell RNA sequencing. Nature immunology. 2016;17(6):666–76.

11. Kanayama M, Izumi Y, Onai N, Akashi T, Hiraoka Y, Ohteki T. Diverse developmental pathways of lymphoid conventional dendritic cells with distinct tissue distribution and function. Science advances. 2025;11(23).

12. Izon D, Rudd K, DeMuth W, Pear WS, Clendenin C, Lindsley RC, et al. A Common Pathway for Dendritic Cell and Early B Cell Development. The Journal of Immunology. 2001;167(3):1387–92.

13. Welner RS, Pelayo R, Nagai Y, Garrett KP, Wuest TR, Carr DJ, et al. Lymphoid precursors are directed to produce dendritic cells as a result of TLR9 ligation during herpes infection. Blood. 2008;112(9):3753–61.

14. Hettinger J, Richards DM, Hansson J, Barra MM, Joschko A-C, Krijgsveld J, et al. Origin of monocytes and macrophages in a committed progenitor. Nature immunology. 2013;14(8):821–30.

15. Yáñez A, Coetzee SG, Olsson A, Muench DE, Berman BP, Hazelett DJ, et al. Granulocyte-Monocyte Progenitors and Monocyte-Dendritic Cell Progenitors Independently Produce Functionally Distinct Monocytes. Immunity. 2017;47(5):890-902.e4.

16. Pronk CJH, Rossi DJ, Månsson R, Attema JL, Norddahl GL, Chan CKF, et al. Elucidation of the Phenotypic, Functional, and Molecular Topography of a Myeloerythroid Progenitor Cell Hierarchy. Cell stem cell. 2007;1(4):428–42.

17. Zhang Y, Gao S, Xia J, Liu F. Hematopoietic Hierarchy - An Updated Roadmap. Trends in cell biology. 2018;28(12):976–86.

18. Lin C-C, Yao C-Y, Hsu Y-C, Hou H-A, Yuan C-T, Li Y-H, et al. Knock-out of Hopx disrupts stemness and quiescence of hematopoietic stem cells in mice. Oncogene. 2020;39(28):5112–23.

19. D’Amico A, Wu L. The Early Progenitors of Mouse Dendritic Cells and Plasmacytoid Predendritic Cells Are within the Bone Marrow Hemopoietic Precursors Expressing Flt3. The Journal of experimental medicine. 2003;198(2):293–303.

